# Allosteric interactions prime androgen receptor dimerization and activation

**DOI:** 10.1101/2022.02.20.481229

**Authors:** Elizabeth V. Wasmuth, Arnaud Vanden Broeck, Justin R. LaClair, Elizabeth A. Hoover, Kayla E. Lawrence, Navid Paknejad, Kyrie Pappas, Doreen Matthies, Biran Wang, Weiran Feng, Philip A. Watson, John C. Zinder, Wouter R. Karthaus, M. Jason de la Cruz, Richard K. Hite, Katia Manova-Todorova, Zhiheng Yu, Susan T. Weintraub, Sebastian Klinge, Charles L. Sawyers

**Affiliations:** Human Oncology and Pathogenesis Program, Memorial Sloan Kettering Cancer Center, New York, NY, 10065 USA; The Rockefeller University, New York, NY, 10065 USA; Structural Biology Program, Memorial Sloan Kettering Cancer Center, New York, NY, 10065 USA; Cryo-Electron Microscopy Facility, Janelia Research Campus, Ashburn, VA, 20147 USA; Molecular Cytology, Memorial Sloan Kettering Cancer Center, New York, NY, 10065 USA; Department of Biochemistry and Structural Biology, University of Texas Health Sciences Center at San Antonio, San Antonio, TX, 78229, USA; Howard Hughes Medical Institute, Memorial Sloan Kettering Cancer Center, New York, NY, 10065 USA

**Keywords:** Nuclear receptor, transcription factors, prostate cancer, allostery, cooperativity

## Abstract

The androgen receptor (AR) is a steroid receptor and master transcription factor that governs gene expression programs required for luminal development of prostate epithelium, formation of muscle tissue and maintenance of the male phenotype. AR misregulation is a hallmark of multiple malignancies, including prostate cancer, where AR hyperactivation and expansion of its transcriptome occur in part through *AR* gene amplification and interaction with oncoprotein cofactors. Despite its biological importance, how AR’s individual domains and its protein cofactors cooperate to bind DNA have remained elusive. Using a combination of reconstitution biochemistry and single particle cryo-electron microscopy (EM), we have isolated three conformational states of AR bound to DNA. We observe that AR forms a non-obligate dimer, with the buried dimer interface utilized by related ancestral nuclear receptors repurposed to facilitate cooperative DNA binding. We identify surfaces bridging AR’s domains responsible for allosteric communication, that are compromised in partial androgen insensitivity syndrome (PAIS), and are reinforced by AR’s oncoprotein cofactor, ERG, and DNA binding site motifs. Finally, we present evidence that this plastic dimer interface for transcriptional activation may have been adopted by AR at the expense of DNA binding. Our work highlights how fine-tuning of AR’s cooperative interactions translate to consequences in development and disease.

## Introduction

AR signaling is a tightly controlled and multifaceted process, regulated through an orchestra of intramolecular and external cues. A better understanding of the rules governing AR activation is of great importance, as multiple pathologies are associated with aberrant AR transcriptional output, including prostate cancer, androgenetic alopecia, and androgen insensitivity syndrome (AIS). That these disorders present with a spectrum of physical and molecular phenotypes (Cancer Genome Atlas Research, 2015; Jeske et al., 2007; La Spada et al., 1991; Lee et al., 2019; McPhaul et al., 1992; Robinson et al., 2015) suggests AR can exist in fully and partly primed states.

A type I nuclear receptor (NR) and member of the 3-ketosteroid receptor (3K-SR) subfamily, AR encodes an approximately 100 kilodalton (kDa) protein with an intrinsically disordered N-terminal domain (NTD), a DNA binding domain (DBD), a flexible Hinge, and a ligand binding domain (LBD) (Fig. S1A) (Weikum et al., 2018). Androgens, including dihydrotestosterone (DHT) and testosterone, bind to AR’s LBD in the cytosol and facilitate AR’s nuclear translocation.

Nuclear AR binds both palindromic and direct repeats of DNA hexamers known as androgen response elements (AREs) to activate its gene expression program, and is further regulated through association with numerous protein cofactors that bind the NTD or LBD through LXXLL and related motifs (Brooke et al., 2008; Weikum et al., 2018). More than most steroid receptors, AR can tolerate higher levels of sequence degeneracy within its ARE, an important feature required for normal development (Adler et al., 1993; Sahu et al., 2014), with 70% of its cistrome comprised of half-sites and up to 99% exhibiting some level of degeneracy (Massie et al., 2007; Wilson et al., 2016; Yu et al., 2010). While many of these sites are not normally associated with active transcription, the overexpression of AR cofactors in prostate cancer are thought to activate expression of pro-proliferative genes at these lower affinity degenerate sites (Chen et al., 2013; Jin et al., 2014; Liu et al., 2017; Mao et al., 2019; Wasmuth et al., 2020).

Despite decades of work, the structural underpinnings of AR regulation conferred by its domains, auxiliary cofactors, and ARE sequence remains unclear. The prevailing view of NR activation comes primarily through work on the distantly related type II NRs, including HNF-4α and PPARγ-RXR, with crystal structures of these multidomain variants revealing cooperative mechanisms of LBD-mediated dimerization to bind DNA (Chandra et al., 2008; Chandra et al., 2013; Chandra et al., 2017). A similar model of constitutive homodimerization through ligand binding is thought to extend to the steroid receptor family, including 3K-SRs and the so-called ancestral steroid receptors (AnSRs), which include the estrogen receptor (ER) family (Greschik et al., 2002; Huang et al., 2018). Intriguingly, a recent study comparing AnSRs to the more evolved 3K-SRs reported that the LBD of the glucocorticoid receptor (GR), a 3K-SR, is not sufficient to dimerize, hypothesizing instead an integral role for the direct repeats within ARE DNA in promoting DBD-mediated dimerization (Hochberg et al., 2020; McKeown et al., 2014). However, AR’s highly degenerate cistrome is inconsistent with a model whereby dimerization relies on canonical ARE repeats (Massie et al., 2007; Wang et al., 2007; Wilson et al., 2016). Current structural studies also support that 3K-SRs may have acquired a mechanism of activation distinct from other NRs, as their LBDs most often crystallize as monomers (He et al., 2004), in contrast to ER (Greschik et al., 2002). Indeed, the few reported dimeric structures of 3K-SRs exhibit variability around their dimerization interfaces, likely indicative of a low affinity interaction (Bledsoe et al., 2002; Nadal et al., 2017; Williams and Sigler, 1998). Whether allosteric surfaces within the 3K-SR LBD contribute to DNA binding remains unclear as structural information has been limited to individual domains (He et al., 2004; Shaffer et al., 2004) or lacks structural features to unambiguously assign DNA or individual domains (Yu et al., 2020).

Greater clarity on the determinants of 3K-SR activation could also be instructive for novel pharmacological intervention, particularly in metastatic prostate cancer where patients inevitably develop resisance to current AR-targeted therapies, including the anti-androgen enzalutamide (ENZ) (Tran et al., 2009; Watson et al., 2015). Yet, historical barriers have impeded progress on this front, including inherent flexibility between the ordered domains, and poor protein solubility and specific activity. We recently developed a means to isolate active multidomain AR and directly demonstrated NTD-dependent autoinhibition of DNA binding (Wasmuth et al., 2020). Driven by the biology of ETS transcription factor translocations in prostate cancer (Cancer Genome Atlas Research, 2015; Chen et al., 2013; Yu et al., 2010), we introduced the oncoprotein ERG into this system and demonstrated ERG is a bonafide AR cofactor, endowed with a LXXLL-like AR interacting motif (AIM) that can reverse NTD autoinhibition through a DNA-independent association with AR’s LBD (Wasmuth et al., 2020). To gain mechanistic insight to the molecular features that govern AR’s dimerization and activation, we have now leveraged this biochemical reconstitution system to trap a DNA bound AR complex with ERG. Using single particle cryo-EM coupled with cross-linking mass-spectrometry, we have discovered that AR exhibits a surprising mode of tunable dimerization, utilizing surfaces important for interdomain allostery for binding to degenerate DNA sequences, which we provide evidence can be reinforced by ERG.

## RESULTS

### ERG chaperones AR to promote DNA binding

To visualize how AR and ERG cooperate to bind DNA at the single molecule level, we imaged recombinant AR and ERG in the presence of duplex ARE DNA by atomic force microscopy (AFM). For these studies, the 52 kDa full-length (FL) ERG protein was used (Fig. S1A), as the AIM-containing ETS domain is not sufficient to promote cooperative DNA binding in a recombinant protein-based DNA binding assay (Fig. S1B), suggesting additional surfaces beyond the AIM interact with AR. Conversely, for these and subsequent structural and biochemical studies using recombinant protein, we assayed a 43 kDa construct of AR lacking its NTD as we and others demonstrated this domain is intrinsically disordered, is not necessary for ERG association and cooperative stimulation, and contributes to N-C autoinhibition in the absence of NTD-cofactor association (He et al., 2001; Schaufele et al., 2005; van Royen et al., 2012; Wasmuth et al., 2020). Under low salt conditions required for AR to bind ARE DNA specifically and with high affinity (Wasmuth et al., 2020), AR extensively aggregated. ERG, in contrast, was more soluble, and remarkably, prevented AR oligomerization through the formation of larger and more globular complexes (Fig. 1A, Fig. S1C).

**Figure 1.**
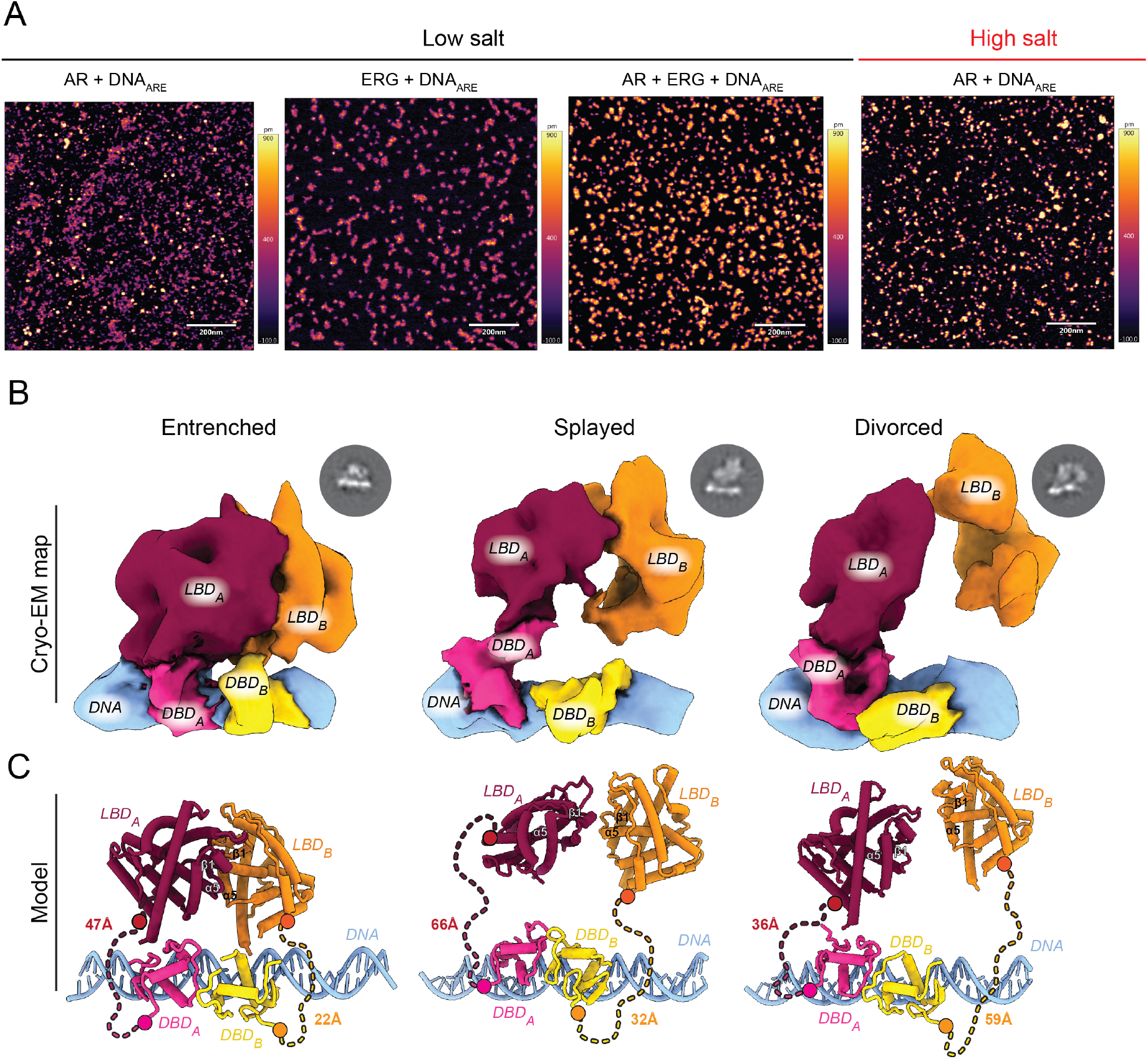
DNA-bound AR exhibits conformational plasticity about its dimer interfaces. (A) Representative AFM images of recombinant AR bound to DNA in the presence or absence of ERG. (B, C) Architecture of three distinct DNA-bound states displaying a spectrum of buried to exposed intermolecular surfaces including the Entrenched, Splayed, and Divorced conformations. (B) Cryo-EM electron density with AR domains and DNA segmented and labeled. Representative 2D classes shown above respective model. (C) Coordinate view derived from X-ray structures of LBD monomers (He et al., 2004) (PBD: 1XOW, red/orange) and the DBD dimer (Shaffer et al., 2004) (PDB: 1R4I, yellow/pink) modeled into cryo-EM maps. Hinge shown as dashed lines, with distance (Å) between the C- and N-termini of the DBD and LBD, respectively, indicated.

### Global architecture of DNA-bound AR

We exploited the dramatic solubilizing effect of ERG to visualize how AR is activated using higher resolution structural methods. We reconstituted and trapped an AR complex designed to model a fully primed state that lacked its NTD, and was bound to DHT, palindromic ARE DNA, and ERG. To enrich for and identify productively bound AR complexes, we performed gentle cross-linking during ultracentrifugation (Stark, 2010) followed by a combination of AFM, negative stain and single particle cryo-EM to screen individual fractions (Fig. S2).

The purified complex was comprised of hetero- and homo-cross-linked species between AR and ERG or with AR, respectively, that migrated around 100 and 150 kDa by SDS-PAGE (Fig. S3A). Using single particle cryo-EM, we isolated three distinct states of AR bound to DNA from this complex mixture that exhibited Entrenched, Splayed and Divorced architectures, with equal number of unique particles among the models (Fig. 1B, C, Fig. S3, Table S1). The resolutions of our structures range from 9.1-11.4 angstroms (Å) with defined features to facilitate docking of X-ray coordinates of individual subunits of AR’s LBD and DBD (Fig. S4) (He et al., 2004; Shaffer et al., 2004). Importantly, the distance between the LBD and DBD of one protomer can be accommodated by the length of the 42 residue disordered Hinge in all three of the models (Fig. 1C).

The most striking difference defining these states is plasticity around a common LBD dimer interface that converges at a prominent surface near beta-sheet 1 and helix 5 of the LBD (Fig. 1, Fig. 2A, Supplemental Video). While all three models share this common dimerization interface, the Entrenched state is the most compact, exhibiting the best fit for the X-ray structure of a LBD dimer (Nadal et al., 2017), whereas the extended conformation in the Divorced state deviates the most (Fig. S5). The fact that AR exists as a non-obligate dimer when bound to DNA is in stark contrast to AnSRs and type II NRs, whose LBDs are sufficient for dimerization at a distinct yet conserved interface (see below) (Hochberg et al., 2020; Weikum et al., 2018).

**Figure 2.**
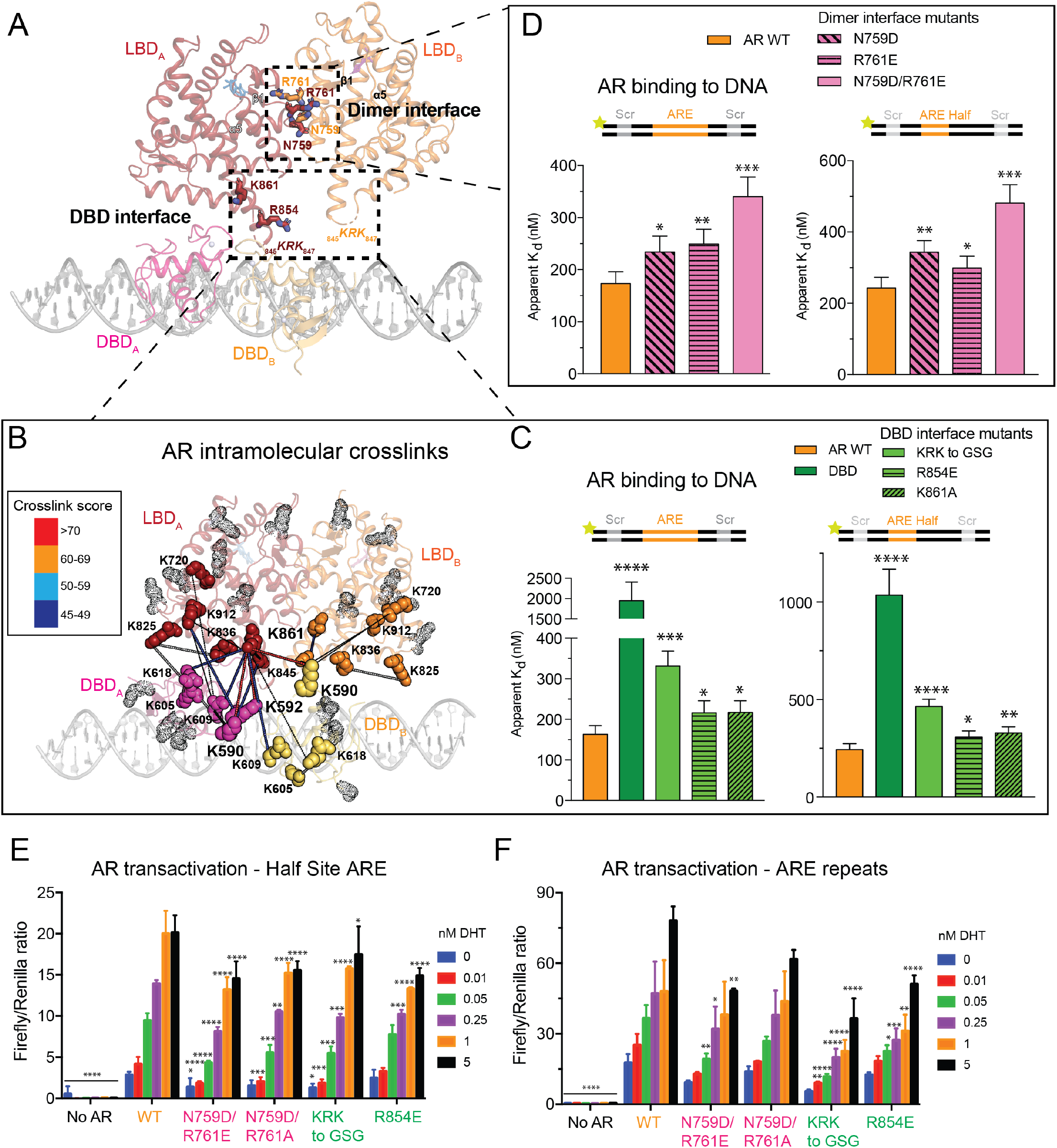
Structural basis for interdomain allostery. (A) Coordinate view of the Entrenched model with LBD residues invoked in the Dimer and DBD interfaces shown as sticks. (B) AR lysines involved in DSSO-mediated intra- and intermolecular cross-links shown as solid spheres on Entrenched model. Rational cross-links are connected by thick lines, and color coded according to score. Lysines not involved in cross-linking are represented as gray transparent dots. (C, D) Fluorescence polarization of AR LBD mutants targeting the (C) DBD- and (D) Dimer interfaces on palindromic (left) and half-site (right) ARE DNA. (E, F) AR transactivation in HEK293 cells on half-site (E) and palindromic (F) ARE reporters.

Density for the two DBDs in all three models are consistent with the head-to-head conformation previously reported in the X-ray structure of the AR DBD dimer bound to non-palindromic ARE repeats (Shaffer et al., 2004). The similar arrangement observed on palindromic (our structure) versus non-palindromic (direct repeat) ARE DNA (Shaffer et al., 2004) could be due in part to flexibility conferred by the Hinge and the conserved five residue “lever arm,” a loop in the DBD previously identified in GR that permits degeneracy within the DNA consensus motif without altering affinity (Meijsing et al., 2009). Reminiscent of the LBD, the DBD dimer interface also displays conformational plasticity, with the DBD of protomer B rotating progressively away from its LBD in the Splayed and Divorced states (Fig. S6). That the DBDs display such flexibility when bound to palindromic ARE DNA suggests DNA sequence is not sufficient to induce uniform dimerization between the DBDs or the LBDs, and that additional surfaces within the LBD likely contribute to cooperative DNA binding.

### Structure-guided mutagenesis reveals interdomain cooperativity

To validate the structural models, we performed cross-linking mass spectrometry (XL-MS) with the lysine cross-linker disuccinimidyl sulfoxide (DSSO) to identify candidate surfaces responsible for interdomain and intermolecular cooperativity (Fig. S7, Table S2). The highest scored cross-links were at the interface between the AR DBD and LBD, independently validating our domain docking and structural observations (Fig. 2B). Interestingly, the two most enriched cross-linked lysines in the DBD, K590 and K592, comprise part of the “lever arm,” which has been speculated to mediate interdomain allostery via its flexibility (Meijsing et al., 2009). Indeed, the “lever arm” lysines primarily cross-link to LBD residues K861 and K847, with these two surfaces in close proximity in both protomers of the Entrenched model, and in one protomer of the Divorced model. K861, the most enriched cross-linked lysine in the LBD, features prominently in the network of interdomain contacts with the DBD. Notably, the equivalent residue in AnSRs is hydrophobic and buried, forming part of the conserved LBD dimerization helix (Fig. S8). In contrast, the 3K-SR family replaced this hydrophobic dimer interface with polar residues, abandoning a constitutive dimerization mechanism for a tunable one (Fig. S8).

We next investigated how LBD contacts at the dimer and DBD interfaces impact AR’s ability to bind DNA by targeting conserved residues identified by our cryo-EM and XL-MS data (Fig. S9, Fig. S10A, B). To benchmark the contributions of the LBD to DNA binding, we also assayed the DBD alone, which bound DNA over fivefold weaker than wild-type (WT) AR (Fig. 2C) and failed to produce DNA supershifts characteristic of WT AR (Fig. S10A), indicating the integral role of the LBD in dimerization and AR activation. Mutation of N759 or R761 at the dimer interface, individually or together, abrogated DNA binding on both ARE palindromic and half site DNA (Fig. 2D), as did perturbation of residues at the DBD interface, including the basic loop KRK 845-847, R854, and K861 (Fig. 2C). Consistent with a role of these LBD residues in cooperative DNA binding, DNA mobility shifts of the mutant proteins were highly altered – the supershifted species in WT were lost in most mutants, instead resembling that of the DBD (Fig. S10A).

To test the model proposed by Thornton and colleagues that ARE repeats within the DNA template drive AR dimerization and transactivation (Hochberg et al., 2020), or whether LBD surfaces contribute to this process, we introduced WT and allosteric mutant AR alleles into cells (Fig. S10C) and measured their ability to activate reporters with either degenerate (half-site) or palindromic ARE sequences (Huang et al., 1999; Zhang et al., 2000). A range of DHT concentrations was used to mimic different levels of AR activity reflective of partially versus fully primed settings. AR transactivation was significantly impaired by both classes of LBD mutations (dimer and DBD interfaces) on the half site reporter, suggesting that the cooperativity conferred through LBD allosteric surfaces enables transactivation of weak AREs (Fig. 2E). Conversely, only the DBD interface mutants showed appreciable transcriptional defects on the palindromic reporter (Fig. 2F), suggesting that when AR is partly primed through limiting ligand or compromised allosteric interactions, DNA consensus repeats can directly promote 3K-SR dimerization (Hochberg et al., 2020; McKeown et al., 2014). Taken together, the impaired DNA binding and transactivation exhibited by the LBD dimer interface mutants on half site DNA (Fig. 2D, E) support a model where direct ARE repeats are not required for AR dimerization.

### ERG impact on differentially primed AR

Interestingly, while ERG is detected in our complexes by XL-MS, SDS-PAGE and immunoblotting (Fig. 3A, Fig. S3A, Fig. S11A, B), we could not resolve features in our structures corresponding to ERG’s PNT and ETS domains. We attribute this in part to the small size of these domains (<15 kDa), which are two to three-fold smaller than the AR LBD and connected by a flexible linker, as well as the resolution of our structural models. Of note, the fact that intramolecular cross-links to AR outnumber intermolecular cross-links to ERG suggests we may have captured states without ERG (Fig. S3A, Fig. S11A, B). Indeed, in the Entrenched model, we observe additional density proximal to the LBDs that can accommodate ERG’s PNT and ETS domains (Fig. S11C, D), consistent with the ERG-AR interface mapped by XL-MS (Fig. 3A, S11A) and our previous work suggesting that ERG interacts with the AR LBD (Wasmuth et al., 2020).

**Figure 3.**
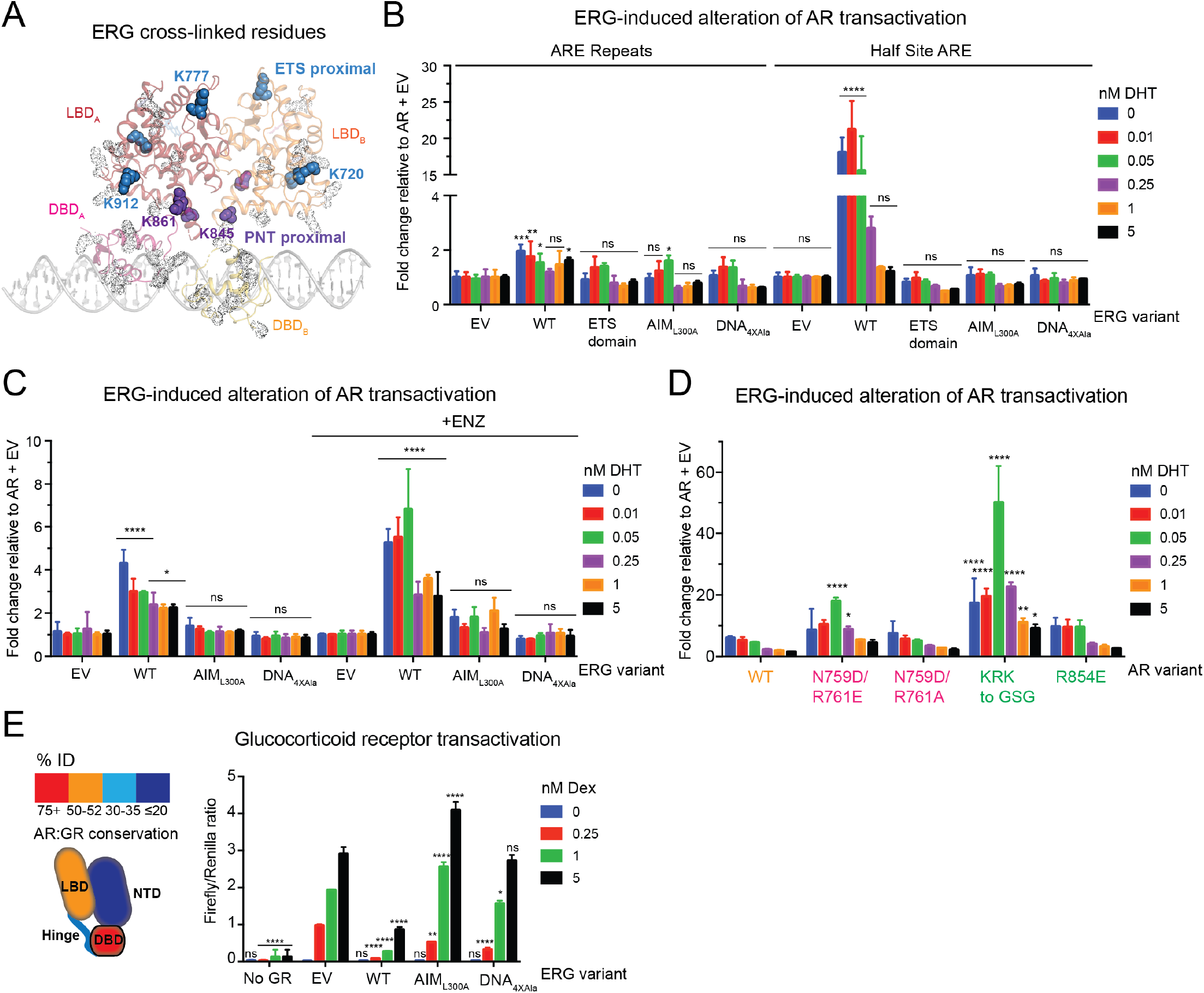
Partly primed AR is more vulnerable to ERG modulation. (A) Intermolecular cross-links between AR and ERG mapped onto Entrenched model. AR lysines that are cross-linked to ERG are shown as solid spheres and colored based on their proximity to ERG’s PNT (purple) or ETS (blue) domains. AR lysines not cross-linked to ERG are represented as gray transparent dots. (B-D) ERG differentially alters AR transactivation (B) of ARE palindromic versus half site reporters, and on the half site reporter (C) in the presence of ENZ, (D) and on AR allosteric mutants. ERG-induced alteration of AR transactivation is calculated by normalizing transactivation of the respective ERG (B, C) or AR variant (D) to the corresponding variant without ERG for a given concentration of DHT. (E) Left: Cartoon depiction comparing sequence conservation between AR and GR. Right: GR transactivation assay with indicated ERG variants with various concentrations of the GR-agonist, dexamethasone (Dex).

One prediction derived from these three conformational states is that the Divorced state, which exhibits the least interdomain connectivity and unassigned electron density (Fig. S11C), may be more susceptible to modulation by cofactors like ERG. To recapitulate what we believe to be the partly primed state of AR based on our finding that ARE repeats can compensate for mutations in the dimer interface (Fig. 2E, F), we measured AR transactivation on half-site versus palindromic ARE repeats and queried whether partly primed AR is more vulnerable to modulation by ERG. Indeed, ERG altered AR transactivation over twenty times more on the ARE half site reporter compared to the palindromic sequence (Fig. 3B). We also performed these experiments in the presence of the anti-androgen, ENZ, which we previously demonstrated allosterically inhibits AR’s ability to bind DNA (Wasmuth et al., 2020), and found that ENZ-inhibited AR was similarly more sensitive to ERG than DHT-activated AR (Fig. 3C).

We next introduced AR and ERG truncations and point mutations in the reporter system to determine the surfaces responsible for these interactions. FL ERG repressed activity of an AR variant lacking the N-terminus, but exerted no effect on AR V7, a splice isoform lacking the LBD that is expressed in CRPC patients and associated with anti-androgen resistance (Watson et al., 2010) (Fig. S11E, F). In contrast, the ERG ETS domain and mutants that fail to interact with AR or with DNA (Wasmuth et al., 2020) largely phenocopied the empty vector (Fig. 3B, Fig. S11E), corroborating the XL-MS data showing surfaces outside the ETS domain interact with the LBD.

We subsequently queried whether the AR LBD mutants were more susceptible to ERG regulation, given that ERG cross-links to AR were detected exclusively along the length of the LBD (Fig 3A, Fig. S11A), and because partly primed AR (as measured by AR half-site activation) is more sensitive to ERG (Fig. 3B, C). Intriguingly, ERG had a pronounced effect on mutant AR transactivation on the half site reporter relative to WT, particularly when DHT concentrations were limiting (Fig. 3D), whereas ERG effects between WT and mutants were virtually indistinguishable on the palindromic reporter (Fig. S11G, H). Similarly, ERG interacted with and enhanced the abilities of the AR LBD mutants to bind DNA (Fig. S11I-K).

Because the LBDs and LXXLL cofactor interacting surfaces of 3K-SRs are highly conserved (Fig. 3E, Fig. S11L, M), we queried whether ERG could modulate other 3K-SRs, including the mineralocorticoid receptor (MR) and GR, as the latter can bypass AR blockade to drive ENZ resistance in CRPC (Arora et al., 2013). Similar to what was observed with AR, WT ERG significantly repressed both GR and MR transactivation, while the AIM and DNA binding mutants were far less potent (Fig. 3E, Fig. S11L). Taken together, our findings show that ERG-dependent modulation of AR activity depends on the LBD through an interaction conserved in other 3K-SRs, and unlike other classes of coactivators, does not require the NTD (Yu et al., 2020).

Having documented selective effects of ERG on AR transactivation on AR half-site reporters, we next turned to a more physiologically relevant model of prostate cancer in which ERG overexpression drives a basal to luminal transition and transcription of a class of AR co-dependent genes whose ARE and ETS binding sites are separated by half a helical turn of DNA in primary mouse prostate organoids lacking *Pten* (Chen et al., 2013; Karthaus et al., 2014; Li et al., 2020; Mao et al., 2019; Wasmuth et al., 2020). Overexpression of WT ERG but not AIM or DNA binding mutants promoted organoid formation as well as expression of luminal marker and AR-ERG co-dependent genes (Fig. S12). Consistent with our model that partially primed AR is more vulnerable to ERG modulation, we noted organoid establishment and gene expression were more pronounced under conditions favoring low AR activity, either through ENZ treatment or DHT withdrawal (Fig. S12B,C).

### The Hinge as a transcriptional tuner

The results presented thus far suggest that allosteric LBD interactions in *cis* with the DBD, or in *trans* either with another LBD or with ERG, can promote AR function. Because the Divorced conformation lacks dimeric contacts between the AR LBDs in contrast to the Splayed and Entrenched conformations (Fig. S11C, D), we queried whether the distances observed between the monomers in the three states correlate with AR’s DNA binding activity (Fig. 1B, C). As a first test of this hypothesis, we engineered fusions of AR monomers separated by 18 or 27 residue linkers to model dimers in either forced or extended proximity, respectively (Fig. S13A, B). We found that the fusion with the shorter of the linkers (AR-AR_18Linker_) bound DNA nearly five-fold tighter than WT AR, while the fusion with the longer linker (AR-AR_27Linker_) bound DNA almost three-fold weaker (Fig. S13D). Thus, the distance between AR protomers critically impacts DNA binding affinity.

Having demonstrated the importance of protomeric spacing, we next postulated that the disordered Hinge connecting the DBD and LBD in part drives these extensive conformational rearrangements (Fig. 1B, C), independent of its intramolecular interactions with AR (Fig. 4A). Curiously, 3K-SR hinges have on average the longest hinge lengths among NR family members, further suggesting a functional role of hinge length in the evolution of this subclass (Fig. 4B). To model the shortest (HNF-4α, a type II NR) and longest (MR) known NR hinges, we altered AR’s 42 residue Hinge by 20 amino acids in regions of uncharacterized function (Clinckemalie et al., 2012) (Fig. S13C). Similar to the findings with the dimeric fusions (Fig. S13D), the short- and long-hinged variants exhibited gain- and loss-of-function in DNA binding relative to WT, respectively, while the short-hinged variant failed to be stimulated by ERG (Fig. 4C, Fig. S11I, Fig. S13E). These results support a model where forced proximity enhances interdomain cooperativity. We next introduced the altered hinge variants into cells and measured their effects on AR transactivation and their ability to be modulated by ERG in the half site reporter assay (Fig. 4D, Fig. S13F-I). Although both hinge variants displayed impaired activity compared to WT AR (consistent with evolutionary selection of an optimal hinge length in cells), we did observe that the short-hinged variant was minimally affected by ERG, consistent with our biochemical observations, whereas ERG induced up to a sixty-fold difference in transactivation in the long-hinged variant compared to WT AR (Fig. 4D, Fig. S13H). Interestingly, hinge length did not impact ERG modulation on the palindromic reporter, consistent with repeat AREs promoting dimerization (Fig. S13G, I) (Hochberg et al., 2020).

**Figure 4.**
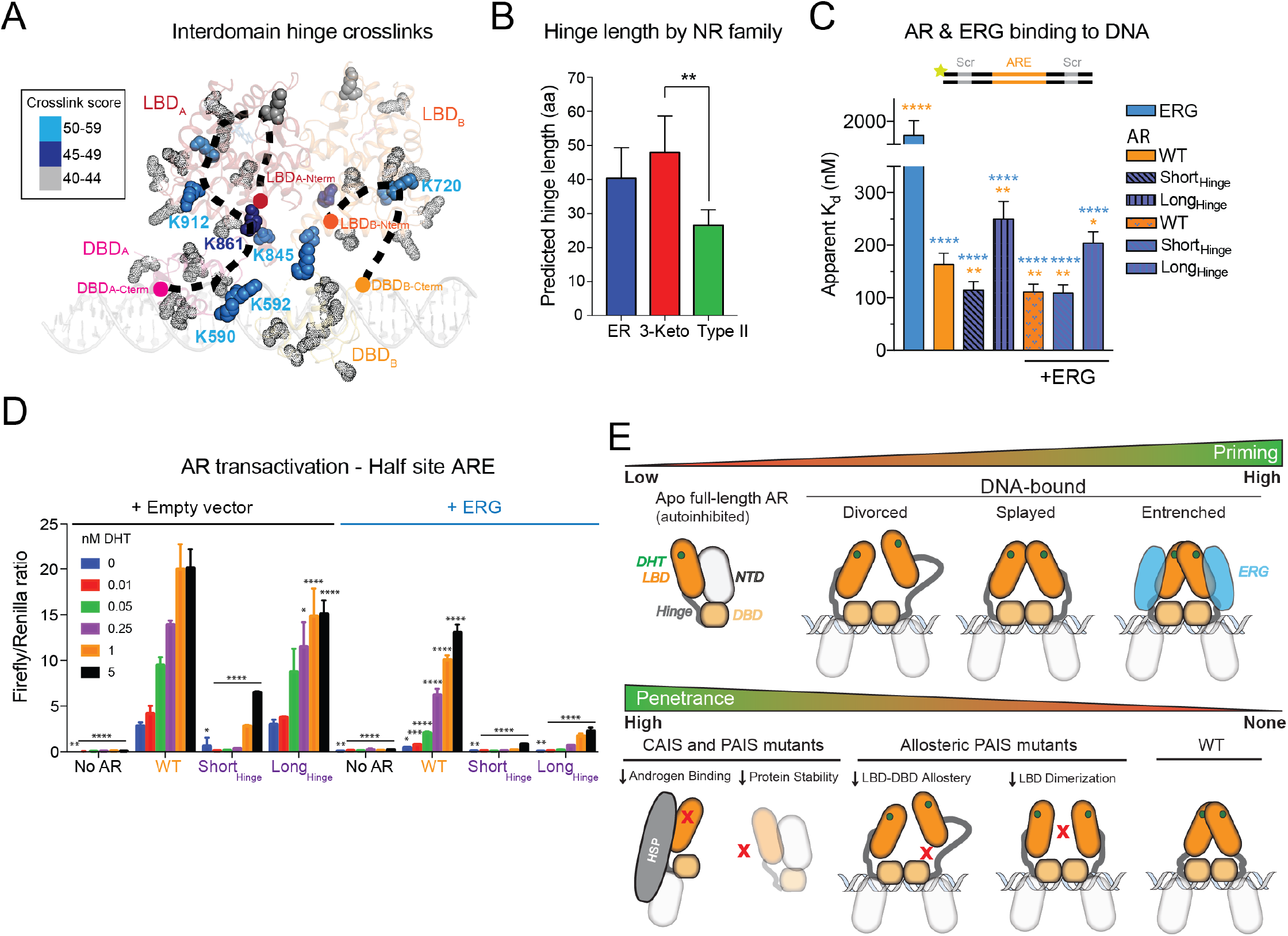
Hinge-mediated flexibility as a transcriptional adaptation. (A) Hinge residues cross-linked to indicated lysines by DSSO are shown as solid spheres, color coded by score, and mapped onto the Entrenched model. (B) Analysis of hinge length across nuclear receptor families. (C) Fluorescence polarization of recombinant AR variants on palindromic ARE DNA in the absence or presence of ERG. (D) AR transactivation with hinge-altered AR in the absence or presence of ERG. (E) Models of AR (top) priming as a function of conformation and ERG status, and (bottom) levels of penetrance caused by complete (CAIS) and partial (PAIS) androgen insensitivity syndrome mutations, including PAIS mutations at surfaces of interdomain allostery.

## DISCUSSION

Regulation of AR is multimodal, with androgen binding, relief of NTD-mediated autoinhibition, and NTD-cofactor interactions contributing to AR priming (He et al., 2001; He et al., 2004; Schaufele et al., 2005; Wasmuth et al., 2020; Yu et al., 2020). While a recent cryo-EM study investigated the role of the NTD in cofactor recruitment (Yu et al., 2020), here we focus on the AR LBD and how this surface can be hijacked by an oncoprotein cofactor. Our study revealed how four additional components directly contribute to AR regulation: 1) LBD allosteric interactions, 2) Hinge length, 3) a full-length LBD-interacting cofactor (ERG), and 4) the composition of the ARE consensus site (Fig. S13J).

We identified two surfaces on the LBD that promote DNA binding and AR transactivation, including intramolecular contacts with the DBD and a plastic dimer interface between two protomers, distinct from the constitutive dimerization interface shared among AnSRs (Fig. S8). We additionally resolved three conformational states of AR bound to DNA representing a continuum of AR activation based on the extent of engagement among these allosteric surfaces (Fig. 4E). Notably, germline mutations within the dimer (N759 and R761) and DBD interfaces (R846 and R854) have been detected in individuals presenting with PAIS, underscoring the physiological importance of these surfaces (Boehmer et al., 2001; Jeske et al., 2007; McPhaul et al., 1992). In contrast to mutations causing complete AIS that are known to cause loss of AR expression, impair androgen binding or are otherwise structurally destabilizing (Boehmer et al., 2001; Chen et al., 2020), these PAIS mutants are examples of how disruption of interdomain allostery translates to more subtle yet pathological consequences (Fig. 4E).

The Hinge also directly contributes to these dynamic states. Our data suggest that this region has expanded in length over evolution to broadly promote transactivation, rather than maintain higher DNA binding affinities through increased steric constraints. In support of this, we noted the gain-of-function conferred by the short hinge was less pronounced on half site compared to full palindromic DNA. Conversely, ERG stimulated the AR variant with a long hinge more on half site DNA (Fig. 4C; Fig. S13E). It remains to be seen whether this adaptation was acquired to facilitate binding to AR’s largely degenerate cistrome *in vivo* (see more below) (Massie et al., 2007; Wilson et al., 2016). *Trans* factors serve to reinforce these interactions, as we have shown for the protein cofactor ERG and the nature of the ARE DNA consensus site. Overall, our data suggest that ERG cooperative interactions more readily influence partly primed AR through interactions with the LBD by potentially inducing a more compact state, providing proof-of-principle evidence for how overexpression of an AR cofactor can confer ENZ resistance (Fig. S12B). Conversely, high affinity ARE repeats rather than half sites or degenerate AREs can directly increase AR binding to DNA and transactivation by promoting dimerization, boosting AR output when AR is impaired through anti-androgens, low DHT, or mutation of its allosteric surfaces.

In summary, AR is hard-wired to tolerate levels of sequence degeneracy for proper development, with the enhanced flexibility endowed by a plastic dimer interface and an extended hinge reported here potentially serving as regulatory mechanisms that allows AR to bind primarily half sites and other degenerate sequences *in vivo* (Massie et al., 2007; Sahu et al., 2014; Wilson et al., 2016; Yu et al., 2010). The transition from Divorced to Entrenched states lends insight to how AR can acquire a broader repertoire of target genes and turn into an oncoprotein under conditions when its allosteric interactions are reinforced. A plastic dimer interface would allow cofactors, such as ERG, to push this conformational equilibrium and fine tune partly primed AR dimers or potential monomers through a graded rheostat mechanism, rather than an on-off switch. Indeed, a dynamic range of AR signaling is observed in prostate cancer cells, independent of AR expression (Lee et al., 2019), with oncogenic cofactors including ERG, reprogramming the AR cistrome and promoting disease progression by redistributing AR binding from higher affinity ARE repeats to lower affinity half sites or degenerate sequences (Chen et al., 2013; Jin et al., 2014; Mao et al., 2019; Norris et al., 2009). This scenario is in contrast to that of the AnSRs that exist as obligate dimers, exhibit less flexibility, and maintain binding to repeat consensus sequences *in vivo* (70%), even in the presence of oncoprotein cofactors (Arruabarrena-Aristorena et al., 2020; Chandra et al., 2008; Chandra et al., 2013; Chandra et al., 2017; Greschik et al., 2002; Huang et al., 2018; Lin et al., 2007). How the multiple conformations and subsequent diversification of AR binding surfaces resulting from this evolutionary adaptation shape the AR regulatory program in development and disease will be a subject of continued investigation.

## LIMITATIONS OF THE STUDY

Although the medium resolution of our structural models allowed us to discover the dynamic nature of DNA-bound AR as well as key allosteric contacts contributing to its DNA binding and transactivation activities, higher resolution cryo-EM structures of ternary complexes between AR, ERG, and DNA are required to elucidate the molecular features responsible for ERG-induced cooperativity. Furthermore, while our previous work (Wasmuth et al., 2020) showed that an AR construct lacking its N-terminus faithfully captures LBD-cofactor regulation subsequent to displacement of the NTD-LBD interaction and engagement wih DNA, future structural studies with full-length AR and ERG are required to discover whether the AR NTD adopts additional regulatory interactions.

## ACKNOWLEDGMENTS

We thank Thomas Walz and Edward Eng for their insights, and Tejasveeta Nadkarni for cloning assistance. Some of this work was performed at the National Center for Cryo-EM Access and Training (NCCAT) and the Simons Electron Microscopy Center located at the New York Structural Biology Center, supported by the NIH Common Fund Transformative High Resolution Cryo-Electron Microscopy program (U24 GM129539) and by grants from the Simons Foundation (SF349247) and NY State Assembly Majority. We acknowledge the Memorial Sloan Kettering Cancer Center Cryo-EM Core Facility funded by NCI Cancer Center Support Grant P30-CA008748. Mass spectrometry analyses were conducted at the University of Texas Health Science Center at San Antonio (UTHSCSA) Institutional Mass Spectrometry Laboratory, with expert technical assistance of Sammy Pardo and Dana Molleur, supported in part by UTHSCSA and the University of Texas System Proteomics Core Network for purchase of the Orbitrap Fusion Lumos mass spectrometer. This research was supported in part by the Department of Defense under award number W81XWH-18-1-0182 (EVW), the Prostate Cancer Foundation Young Investigator Award (EVW), the National Institute of General Medical Sciences of the National Institutes of Health under award number K99 GM140264 (EVW), the Howard Hughes Medical Institute (CLS) and the National Cancer Institute (CLS). CLS is an investigator of the Howard Hughes Medical Institute. The content is solely the responsibility of the authors and does not necessarily represent the official views of the National Institutes of Health.

## AUTHOR CONTRIBUTIONS

EVW, AVB, WRK, NP, JCZ, STW, SK, and CLS designed experiments. EVW, JRL, EAH, KEL, KJP, DM, BW, and MJDLC performed experiments. WF and PAW contributed reagents. Cryo-EM data were analyzed by EVW, AVB, NP, RKH and SK. Negative stain EM data were analyzed by EVW, DM, and ZY. AFM data were acquired by BW and analyzed by BW and KMT. XL-MS data were analyzed by STW. EVW, AVB, SK, and CLS interpreted results. EVW and CLS wrote the manuscript. All authors contributed to the manuscript.

## ADDITIONAL INFORMATION

EM maps are deposited on EMDB with accession codes EMD-25132, EMD-25133 and EMD-25134.

## METHODS

### Recombinant protein expression and purification

Recombinant mouse AR lacking the N-terminus and human ERG proteins were cloned, expressed and purified as described previously (Wasmuth et al., 2020). All constructs contained N-terminal Smt3 fusions. A N-terminal truncation of human estrogen receptor (ER) alpha isoform 1 corresponding to amino acids 176-595 was codon optimized for expression in *E. coli* (Genscript) and subsequently cloned into pRSF-Duet1 (Novagen). The ETS domain of human ERG isoform 2 (residues Q272-E388) was cloned into pRSF-Duet1. The AR DBD (residues D548-E651) was cloned into pET-Duet1 (Novagen). All AR mutants were cloned by HiFi assembly (NEB). For AR Hinge variants, residues of unknown function proximal to the LBD (652-671) were excised, as this region is distal from the bipartite nuclear localization sequence previously implicated in DNA binding and acetylation (Haelens et al., 2007). For AR Long-Hinge, a 20 residue Gly-Ser linker was introduced between residues 651 and 652. Briefly, all expression plasmids were transformed into BL21DE3 codon plus cells (Novagen) and protein expression induced by addition of 0.1mM isopropyl-β-D-thiogalactoside and overnight shaking at 16°C. Cells were lysed by French press, and supernatants purified by Ni-NTA (Qiagen), followed by affinity purification on heparin Hi-Trap (Cytiva Life Sciences), overnight cleavage of the Smt3 tag by Ulp1, and final purification by size exclusion chromatography on either Superdex 200 or Superdex 75 (Cytiva Life Sciences) in a final buffer of 350 mM NaCl, 40 mM HEPES pH7.5, 1 mM TCEP [tris(2-carboxyethyl)phosphine] for ERG proteins, 350 mM NaCl, 40 mM HEPES pH7.5, 1 mM TCEP, 5% glycerol and 20 μM DHT for AR constructs, and 350 mM NaCl, 40 mM HEPES pH7.5, 1 mM TCEP, 10% glycerol and 20 μM beta-estradiol for ER.

### Protein cross-linking

AR, ERG and the indicated ARE DNA were mixed to a final concentration of 10 μM for 1 hour on ice and then dialyzed to 150 mM NaCl, 40 mM HEPES pH7.5, 1 mM TCEP, 20 μM DHT, 0.01% NP40. For structural characterization by AFM and electron microscopy, complexes were then subjected to Grafix (Stark, 2010) to cross-link and simultaneously separated by size. Individual fractions were quenched with 50 mM Tris-HCl pH 8.0 after ultracentrifugation. For mass spectrometry, the reconstituted AR, ERG and ARE_35_ DNA complex was cross-linked with 800 μM DSSO for 1 hour on ice before being quenched with 50 mM Tris-HCl pH 8.0 and then separated by ultracentrifugation. Fraction 11 of the Grafix and DSSO cross-linked samples were further analyzed by cryo-EM and mass-spectrometry, respectively.

### DNA binding assays

Unlabeled and 5’ fluorescein-labeled duplex DNAs were purchased from IDT and had the following sequences, with ARE sites in bold and ETS sites in italics: ARE/Scr: 5’ TACCTAGCGTGGCC**AGAACA**TCA**TGTTCT**CCGGTGCGATCCAG 3’; ARE/ETS 6bp: 5’ TACC*GGAA*GTGGCC**AGAACA**TCA**TGTTCT**CCGGTG*AAGG*CCAG 3’; ARE-Half-Site/Scr: 5’ AGACCTAGCGTGGCC**AGAACA**TCATTAAGCCCGGTGCGATCCAG 3’; ARE_25_: TGGCC**AGAACA**TCA**TGTTCT**CCGGT; ARE_35_:

TAGCGTGGCC**AGAACA**TCA**TGTTCT**CCGGTGCGAT. Binding buffer consisted of 150 mM NaCl, 40 mM Tris pH8.0, 1 mM TCEP, 20 μM DHT, 10% glycerol, 0.01% NP40. As described previously (Wasmuth et al., 2020), equimolar amounts of AR and ERG were pre-incubated on ice for 30 minutes before mixing with the specified dsDNA. For DNA gel shift assays, 50 nM of unlabeled DNA was incubated with 250 nM of total protein on ice for 1 hour. Gel shifted products were resolved on 4-20% TBE PAGE and DNA stained with Sybr Gold (Thermofisher Scientific). For fluorescence polarization experiments measuring DNA binding, 100 nM fluorescein-labeled dsDNA was incubated for 30 minutes on ice with increasing concentrations of the indicated protein (0 to 4 μM final concentration). Data from triplicate experiments was analyzed, and when applicable a model for receptor depletion was used to calculate apparent K_d_ values with Prism, GraphPad Software. Data were analyzed using one-way ANOVA, with ****P<0.0001; n.s. (not significant), P>0.05. Data presented as mean +/− standard deviation from n=4 experiments.

### Atomic force microscopy

Protein-DNA complexes were prepared as described in previous section as native (Main Figure 1, Figure S1) or Grafix cross-linked forms (Figure S2). Native complexes were diluted to 50 nM in low salt (DNA binding buffer) or high salt buffer immediately before imaging. 20 μl of sample was applied to a freshly cleaved mica and rinsed with ultrapure deionized water twice before being gently dried with UHP argon gas. An Asylum Research MFP-3D-BIO (Oxford Instruments, Goleta CA) was used to image in tapping mode. The samples were imaged in air, at room temperature, and under controlled humidity. A silicon nitride probe Olympus AC240 (Asylum Research, Goleta CA) with resonance frequencies of approximately 70 kHz and spring constant of approximately 1.7 N/m was used for imaging. Images were collected at a speed of 1 Hz with an image size of 1 μm at 512 × 512-pixel resolution. Raw data were exported into 8-bit grayscale Tiff images using the Asylum Research’s Igor Pro software and imported into FIJI/ImageJ (NIH) for quantification of volume in Figure S2 using a custom written FIJI code.

### Negative stain electron microscopy

3 microliters of the indicated Grafix fraction with ARE_35_ DNA were applied on glow discharged 400 mesh copper grids with carbon support (Electron Microscopy Sciences) and stained with Nano-W (Nanoprobes). Datasets were collected at 120 kV on a Tecnai T12 microscope (Thermofisher Scientific) and consisted of 264 micrographs and 107,445 particles (Fraction 9) and 365 micrographs and 61,034 particles (Fraction 14). Data processing, including particle autopicking and 2D classification were performed in cisTEM (Grant et al., 2018).

### Cryo-electron microscopy

3.5 microliters of Grafix purified complex with ARE_35_ DNA (Fraction 11) was applied to glow discharged R1.2/1.3 holey carbon grids (Quantifoil) at 4ºC and plunge frozen in liquid ethane using a Vitrobot Mark IV (Thermofisher Scientific). Data were collected at 300 kV on a Titan Krios (Thermofisher Scientific) with energy filter using a K3 Summit Detector in counting mode. 17,798 images were recorded at 1.069Å per pixel with a nominal magnification of 81,000. A total dose of 61.27 e^-^/Å was fractionated over 50 frames, with a defocus range of −0.8 μm to −2.5 μm. Frames were motion-corrected using MotionCor2 (Zheng et al., 2017) and the contrast transfer function estimated using CTFFIND4 (Rohou and Grigorieff, 2015). Particles were picked with crYOLO (Wagner et al., 2019) and subsequently 2X binned. A 3D ab initio model was first obtained in Cryosparc (Punjani et al., 2017), and then imported to RELION-3 (Zivanov et al., 2018) for subsequent rounds of 3D classification and refinement of the entire dataset. The final Entrenched, Splayed, and Divorced reconstructions consist of 68,581, 53,169, and 51,454 particles, with resolutions of 11.4, 9.1, and 9.4 Å, respectively. ARE_35_ DNA was modeled in Coot (Emsley et al., 2010). Individual domains of the AR LBD and DBD were manually docked into respective EM maps and subject to rigid body refinement in Chimera (Pettersen et al., 2004) using the PDB coordinates for the AR LBD (1XOW) (He et al., 2004) and DBD (1R4I) (Shaffer et al., 2004). Composite PDBs of the ∆NTD AR dimer bound to DNA showed good fit into corresponding cryo-EM density with correlation coefficients of 0.8318, 0.8267, and 0.8717, for the Entrenched, Splayed, and Divorced models, respectively. Segmentation of individual domains of AR was performed using Segger (Pintilie et al., 2010) in ChimeraX (Pettersen et al., 2021). Figures were rendered in ChimeraX and PyMol.

### Immunoblotting

For detection of recombinant proteins, 2 μl of Grafix and DSSO fractions were diluted 1:20 and run on 4-12% SDS-PAGE, transferred to PDVF, and detected by ECL Prime (Cytiva Life Sciences) using HRP-anti-rabbit IgG. For detection of protein from mammalian cells, total protein was extracted by MPER lysis (Thermofisher Scientific), quantitated by BCA assay (Pierce), and 10 μg lysate resolved by 4-12% SDS-PAGE. The following antibodies were used: androgen receptor antibody (Abcam ab52615 - 1:1000 for lysates, 1:5000 for recombinant protein), ERG (Abcam ab92513 - 1:1000 for lysates, 1:2000 for recombinant protein), B Actin (Cell Signaling 4970S - 1:5000,) and Cyclophilin B (Cell Signaling 43603S - 1:5000).

### Cross-linking mass-spectrometry

DSSO cross-linked complexes were separated by SDS-PAGE (12% Bis-Tris) and stained with Coomassie blue. The bands of interest (~100 kDa) were manually excised, individually reduced in situ with TCEP and alkylated in the dark with iodoacetamide prior to treatment with trypsin (Promega, sequencing grade). Each digest was analyzed by capillary HPLC-electrospray ionization tandem mass spectrometry on a Thermo Scientific Orbitrap Fusion Lumos mass spectrometer. On-line HPLC separation was accomplished with an RSLC NANO HPLC system (Thermo Scientific/Dionex): column, PicoFrit (New Objective; 75 μm i.d.) packed to 15 cm with C18 adsorbent (Vydac; 218MS 5 μm, 300 Å). Precursor ions were acquired in the orbitrap in centroid mode at 120,000 resolution (*m/z* 200); data-dependent higher-energy collisional dissociation (HCD) spectra were acquired at the same time in the linear trap using the “top speed” option (30% normalized collision energy). Other MS scan parameters included: mass window for precursor ion selection, 0.7; charge states, 2 – 5; dynamic exclusion, 15 sec (± 10 ppm); intensity to trigger MS^2^, 50,000. Mascot (v2.7.0; Matrix Science) was used to search the spectra against a combination of the SwissProt database [SwissProt 2019_10 (561,356 sequences; 201,858,328 residues] plus a local database that includes the sequences of the target proteins (578 sequences; 213,622 residues). Cysteine carbamidomethylation was set as a fixed modification and methionine oxidation was considered as a variable modification. Trypsin was specified as the proteolytic enzyme, with two missed cleavages allowed. The Mascot cross-linking feature for DSSO was used for the corresponding sample searches. The results were exported in xiView-CSV format and as a FASTA file containing the identified peptides sequences for import into xiView for data visualization. Cross-linked lysines were depicted in 2D using xiNET cross-link viewer (Combe et al., 2015) and rendered in 3D as solid-colored spheres mapped onto the Entrenched model using PyMol.

### Mammalian construct generation

For transfection-based reporter assays, human GR and MR were cloned into pACT (Promega) as N-terminal VP16 fusions using HiFi assembly (NEB). The human ERG ETS domain, and human VP16-AR WT and mutants were cloned into pCDNA3.1 using HiFi assembly. All other constructs were described previously (Wasmuth et al., 2020). For lentiviral transduction of ERG variants in mouse prostate organoids, modified derivatives of pLVX-TRE3G-IRES (Takara) were engineered for constitutive expression by replacing the TRE3G element with a UBC promoter. To construct LVX-eGFP-ERG-PuroR variants, eGFP was cloned into MCS I; human ERG variants were cloned via HiFi assembly into MCS II.

### Cell culture

HEK293T cells were cultured in DMEM with high glucose, 10% fetal bovine serum, and penicillin streptomycin. Mouse prostate organoids were isolated and cultured in Matrigel (Corning) using standard methods (Karthaus et al., 2014). Pten^−/−^;sgERG organoids previously generated by electroporation of Cas9-sgRNA ribonucleoprotein complexes against ERG (Feng et al., 2021) were transduced with an empty vector control, or an allelic series of WT and mutant ERG variants and selected with 2 μg/mL puromycin for 1 week. Cells were cultured in standard mouse prostate organoid media, supplemented with 5 ng/mL EGF and 1 nM DHT. Pten^−/−^;ERG organoids were confirmed to be GFP positive throughout the duration of the experiment by fluorescence microscopy. All lines were confirmed to be free of mycoplasma using the Lonza MycoAlert Mycoplasma Detection Kit (LT07-318).

### AR reporter assays

The ARE repeat reporter (4X-ARE, firefly luciferase) has been previously described (Tran et al., 2009). The half site ARE reporter is derived from the minimal rat probasin sequence (Zhang et al., 2000) and was cloned into pGL3 via KpnI and NcoI restriction sites (Promega). pRL-TK (Promega) was used as a Renilla luciferase normalization control. Assays were performed as described previously (Wasmuth et al., 2020). Briefly, plasmids were transfected into HEK293 cells in triplicate using Lipofectamine 2000 (Thermofisher Scientific) in the presence of various amounts of DHT, and luciferase activity read 36 hours after transfection using Dual Glo reagent (Promega). To calculate overall AR transactivation, firefly luciferase activity was normalized to Renilla, and is represented as Firefly/Renilla ratio. ERG-induced alteration of AR transactivation is the fold change between AR transactivation in the absence (empty vector) and presence of WT ERG for a given AR variant. Data were analyzed using two-way ANOVA, with ****P<0.0001; n.s. (not significant), P>0.05. Data presented as mean +/− standard deviation from n=3 experiments.

### Organoid establishment assays

Pten^−/−^;ERG organoids were trypsinized to single cells and plated at a density of 84 cells per 25 μl dome of Matrigel, during which time EGF was removed from the media. Cells were treated with either no ligand (0 nM DHT), with DHT (1 nM), or with enzalutamide (10 μM) for 11 days before quantifying establishment, refreshing media every 2-3 days. Percent formation was derived from number of established organoids divided by the total number of cells plated, multiplied by 100. Data from an average of 8 replicate wells were analyzed using two-way ANOVA, with ****P<0.0001; n.s. (not significant), P>0.05. Data presented as mean +/− standard deviation from n=3 experiments.

### RNA extraction and quantitative PCR (qPCR)

RNA was extracted from organoids using an RNeasy Kit (Qiagen) followed by on-column DNase treatment. cDNA was generated with the High Capacity cDNA Reverse Transcription Kit (Thermofisher Scientific). qPCR was performed with QuantiFast SYBR Green PCR master mix (Qiagen). Primers used for mouse Actb (QT00095242), Ar (PPM05196F), Krt8 (PPM04776F), Nkx3-1 (PPM05232A), and Tmprss2 (QT00156093) were purchased from Qiagen. Data were analyzed using the ∆Ct method and statistical significance calculated using two-way ANOVA, with ****P<0.0001; n.s. (not significant), P>0.05. Data presented as mean +/− standard deviation from n=3 experiments.

## SUPPLMENTAL INFORMATION

Figures S1-13.

Tables S1-2.

Supplementary Movie. Plasticity of the AR dimer interface is apparent in the transition of Entrenched to Divorced states.

